# Testing pseudo-linear models of responses to natural scenes in primate retina

**DOI:** 10.1101/045336

**Authors:** Alexander Heitman, Nora Brackbill, Martin Greschner, Alexander Sher, Alan M. Litke, E.J. Chichilnisky

## Abstract

A central goal of systems neuroscience is to develop accurate quantitative models of how neural circuits process information. Prevalent models of light response in retinal ganglion cells (RGCs) usually begin with linear filtering over space and time, which reduces the high-dimensional visual stimulus to a simpler and more tractable scalar function of time that in turn determines the model output. Although these pseudo-linear models can accurately replicate RGC responses to stochastic stimuli, it is unclear whether the strong linearity assumption captures the function of the retina in the natural environment. This paper tests how accurately one pseudo-linear model, the generalized linear model (GLM), explains the responses of primate RGCs to naturalistic visual stimuli. Light responses from macaque RGCs were obtained using large-scale multi-electrode recordings, and two major cell types, ON and OFF parasol, were examined. Visual stimuli consisted of images of natural environments with simulated saccadic and fixational eye movements. The GLM accurately reproduced RGC responses to white noise stimuli, as observed previously, but did not generalize to predict RGC responses to naturalistic stimuli. It also failed to capture RGC responses when fitted and tested with naturalistic stimuli alone. Fitted scalar nonlinearities before and after the linear filtering stage were insufficient to correct the failures. These findings suggest that retinal signaling under natural conditions cannot be captured by models that begin with linear filtering, and emphasize the importance of additional spatial nonlinearities, gain control, and/or peripheral effects in the first stage of visual processing.

## Introduction

Decades of research on the retina have yielded many computational models of its function, starting with the linear description of spatial receptive fields of retinal ganglion cells (RGCs) [1,2]. A central goal of these computational models is to synthesize current understanding and provide quantitative predictions of responses to a range of visual stimuli. Most models of RGC light response begin with a linear filter (the spatiotemporal receptive field) which reduces the high-dimensional spatiotemporal input stimulus to a scalar function of time, enormously simplifying the modeling problem. Subsequent elements in the model then generate a sequence of spikes from the scalar output of the linear filter. Despite their simplicity, these pseudo-linear models can accurately replicate the temporal structure of RGC light responses to stochastic visual stimuli that are commonly used to characterize neural responses in the retina [3–5] (see [6]), suggesting that important aspects of neural computation in the retinal circuitry are approximately linear. These findings validate the concept of a spatiotemporal receptive field for RGCs, which is effectively tied to the linearity assumption. They also permit a compact and powerful summary of neural computation at the earliest stage of visual processing. Indeed, most models of visual computation in the brain begin with the assumption, explicit or implicit, that retinal processing is largely linear.

However, responses to artificial stochastic stimuli may not reveal how the retina functions in the life of the organism. Naturalistic stimuli exhibit distinctive statistics, and numerous studies support the notion that the visual system is specifically tuned to represent stimuli with natural statistics [7–9]. Furthermore, many studies have revealed nonlinear computations, including nonlinear summation over space and time, in several types of RGCs in several species (e.g. [10–16]; see [17]) as well as some of the underlying mechanisms [18–21], using tailored artificial stimuli. These findings are inconsistent with pseudo-linear models of retinal function. However, it is unclear how relevant these nonlinearities are for retinal encoding of natural scenes. It has been suggested that they may not be very significant in the mouse retina [22], though no tests have been performed in the primate retina. In principle, retinal computations could behave mostly linearly in natural conditions, even though nonlinear behavior can be elicited with certain stimuli. Given these diverse findings, a basic question arises: is the first stage of visual processing effectively linear in a natural setting?

The goal of the present study is to answer this question in the retina of the macaque monkey, the animal model with retina, visual system, and visual behaviors most similar to those of humans. To test the adequacy of pseudo-linear models, a natural focus is the *generalized linear model* (GLM), which is among the most recent widely-used models and has several key conceptual and technical advantages over other pseudo-linear models [5]. First, the GLM includes an instantaneous nonlinearity after the linear filter, which captures the nonlinear intensity-response relationship of RGCs. Second, the GLM includes post-spike feedback, which accounts for non-Poisson structure in RGC spike trains, such as bursting and refractoriness, that are not captured by simpler models [23]. Third, the GLM can be extended easily, using post-spike cross-coupling filters, to capture the stimulus-independent correlations present in RGC populations. Fourth, the GLM can be robustly fitted to experimental data, including data from entire populations of coupled cells, because the log-likelihood function is convex in the model parameters [24]. Finally, the GLM permits optimal reconstruction of the stimulus from spike trains, which reveals how specific aspects of the neural response encode the stimulus [25,26]. These features of the GLM make it useful for the study of other neural systems as well [27,28].

Here we test the ability of the GLM to account for RGC responses to movies composed of natural images with simulated fixational eye movements. Using large-scale multi-electrode recordings from macaque retina, we examined responses to these naturalistic stimuli in populations of ON and OFF parasol cells, which initiate the magnocellular pathway of the primate visual system. The results indicate that the performance of the GLM in reproducing responses to naturalistic stimuli is markedly inferior to its previously reported accurate performance with white noise stimuli. We further demonstrate that a GLM modified with pointwise scalar nonlinearities before and after the linear filter also fails to reproduce responses to naturalistic stimuli. These findings indicate that pseudo-linear models are insufficient for explaining the visual signaling function of primate RGCs in natural viewing conditions. They also highlight the importance of building computational models which incorporate known properties of retinal function, including nonlinear spatial summation, gain control, and periphery effects, to understand natural vision.

## Results

### Primate retinal ganglion cell responses to natural scenes

To understand the pattern of retinal responses elicited by natural scenes, large-scale multi-electrode recordings were obtained from retinal ganglion cells (RGCs) in the isolated macaque retina [29,30]. Spiking responses of hundreds of RGCs were examined, and spatiotemporal white noise was used to probe their light responses and classify distinct cell types [30–32]. Complete populations of ON and OFF parasol RGCs covering the region recorded were examined further. Images of natural scenes from the van Hateren database [33] were presented and jittered over time in a manner that simulated fixational eye movements, and were replaced once per second to emulate the rapid transitions that occur at saccades (see Methods, Fig. 1A,B).

**Figure 1.**
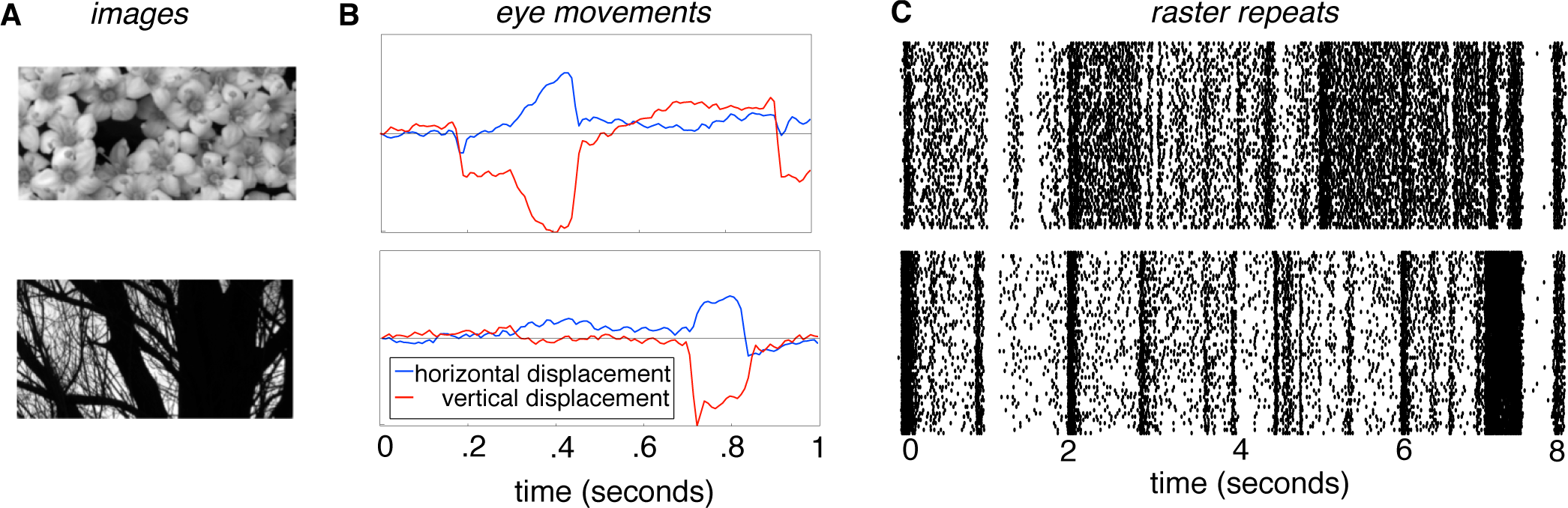
Naturalistic stimuli and responses A. Sample images taken from the Van Hateren database [33]. B. Two one second fixational eye movement traces taken from recordings of awake fixating macaques (see Methods). Note presence of microsaccades, drift and jitter. C. Sample of recorded responses to a natural scenes stimulus (top: ON parasol, bottom: OFF parasol). Each tick represents the occurrence of a spike, each row represents a single presentation of the stimulus (total of 57 presentations). Note strong sustained periods of spiking and silence.

Unsurprisingly, natural scenes evoked patterns of neural response very different from those evoked by spatiotemporal white noise. The combination of natural image structure and emulated eye movements produced strong and extended periods of firing and silence (Fig. 1C), compared to the briskly but weakly modulated responses observed with white noise (Fig. 2C,D,G,H). Thus, the different statistics of these stimuli give rise to different patterns of activity in the retinal output.

**Figure 2.**
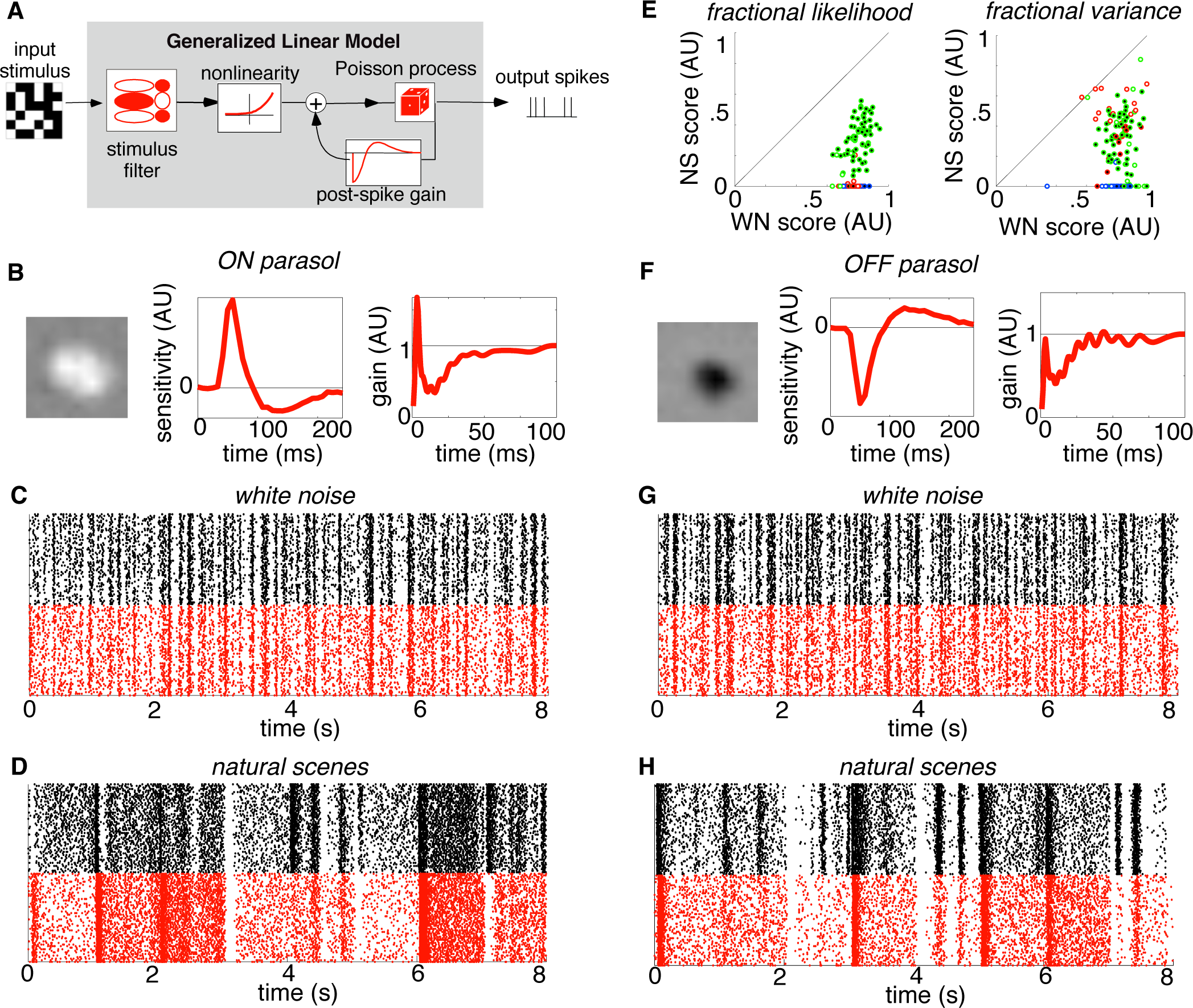
Generalized linear model (GLM) fitted to white noise does not capture responses to natural scenes. A. Schematic of GLM, with linear stimulus filter, response nonlinearity, stochastic spike generation, and post-spike feedback (see Results). B. Sample GLM components obtained from fits to white noise data, for an ON parasol cell. The stimulus filter is decomposed into its separable spatial (left) and temporal (middle) components. The feedback filter (right) is expressed in terms of gain after the exponential nonlinearity (see Methods). C. White noise raster for the same ON parasol cell (black, 57 repetitions). Response structure is approximately reproduced by the fitted model (red). D. Natural scenes rasters for the same ON parasol cell (black, 57 repetitions). Response structure is not reproduced by the GLM fitted to white noise (red). E. Comparison of GLM predictions of responses to white noise (C,G) and natural scenes (D,H), using fractional log-likelihood increment (left) and fraction of explainable variance (right) measures (see Methods). In both cases the GLM was fitted to white noise data. Colors of points denote different retinas. Negative scores were set to zero. White-center points represent ON parasol cells, black-center points represent OFF parasol cells. F-H. Same as B-D for an OFF parasol cell.

### Testing the extensibility of the generalized linear model to natural scenes

To test whether pseudo-linear models can generalize easily to capture the natural function of the retina, a generalized linear model (GLM) was fitted to data obtained with white noise stimuli, and the fitted model was then used to predict light responses to both white noise and natural scenes [5]. In the GLM, the stimulus is first convolved with a spatiotemporal linear filter to produce a univariate generator signal over time. The generator signal drives spike generation through an exponential nonlinearity that controls the rate of an inhomogeneous Poisson point process (Fig. 2A). These portions of the model are designed to capture the integration of light inputs over space and time, the nonlinear relation between stimulus strength and firing probability, and the stochastic nature of retinal responses. In addition, a fitted feedback waveform is added to the generator signal after each generated spike. This term permits the model to capture non-Poisson structure observed in real spike trains, such as refractoriness and bursting [23,34].

Fits of the model parameters to white noise data revealed spatiotemporal filters and feedback filters similar to those seen in previous work (Fig. 2B,2F). With these fitted parameters, the GLM provided an accurate reproduction of the structure of RGC firing to held-out data consisting of repeated presentations of a white noise stimulus, consistent with previous findings (Fig. 2C,2G) [5]. However, this same model, with the same parameters, largely failed to reproduce the distinctive firing structure observed with natural scenes (Fig. 2D,2H).

The inferiority of the model predictions with natural scenes compared to white noise was s consistent in 46 ON and 87 OFF parasol cells examined in three retinas (Fig. 2E). To summarize model performance across many cells, the quality of the model predictions was quantified in two ways (see Methods) that allow for the fact that firing structure is very different in the two stimulus conditions. First the likelihood of the rasters given the fitted model was quantified by the *fractional log-likelihood increment*. This metric is a monotonic transform of likelihood, normalized to [0,1] in a manner that focuses on the model’s ability to capture stimulus-driven response variation over time (see Methods). Second, the ability of the model to directly replicate the response was quantified by the *fraction of explainable variance*. This measure is also normalized to [0,1], and focuses on firing modulations which were reproducible across trials (see Methods).

Using both measures, model prediction accuracy was substantially lower for natural scenes stimuli than for white noise stimuli (Fig. 2E). Note that the absolute model performance differed somewhat in the three retinas recorded, presumably due to differences in the preparations. These differences could include the degree of photopigment bleaching, receptive field sizes relative to stimulus pixels, and other sources of variability between animals and experimental preparations. In all cases, however, the model captured responses to natural scenes much less accurately than responses to white noise. Thus, a GLM fitted to white noise data cannot generalize to explain responses to natural stimuli.

### Testing the structure of the model with natural scenes

One reason for the failure of the GLM to generalize, despite capturing responses to white noise accurately, could be that retinal processing adjusts to the statistics of the stimulus by changing its properties (see [35]). Adaptive changes in retinal processing could potentially be captured by allowing the parameters of the GLM to change in different stimulus conditions. If this were true, the GLM would remain useful for summarizing retinal responses, with the understanding that model predictions are accurate only if the parameters of the GLM are estimated using stimuli with similar statistics, because of adaptation in the retina.

To test this possibility, the GLM was fitted to data obtained with a subset of natural scenes stimuli, and then used to predict the responses to a different subset of natural scenes stimuli (cross-validated). This approach produced somewhat more accurate predictions of light responses (Fig. 3C,3F). These improvements took the form of both minor corrections in the structure of the response as well as overall changes in firing rate. However, the discrepancies between data and model remained large compared to those obtained with white noise (Fig. 3D). Across three preparations, the results were consistent: fitting and testing the model with natural scenes produced substantially less accurate predictions of response than fitting and testing with white noise (Fig. 3D). Thus, the parametric form of the GLM is less well-suited to capturing light responses with natural stimuli than with white noise.

**Figure 3.**
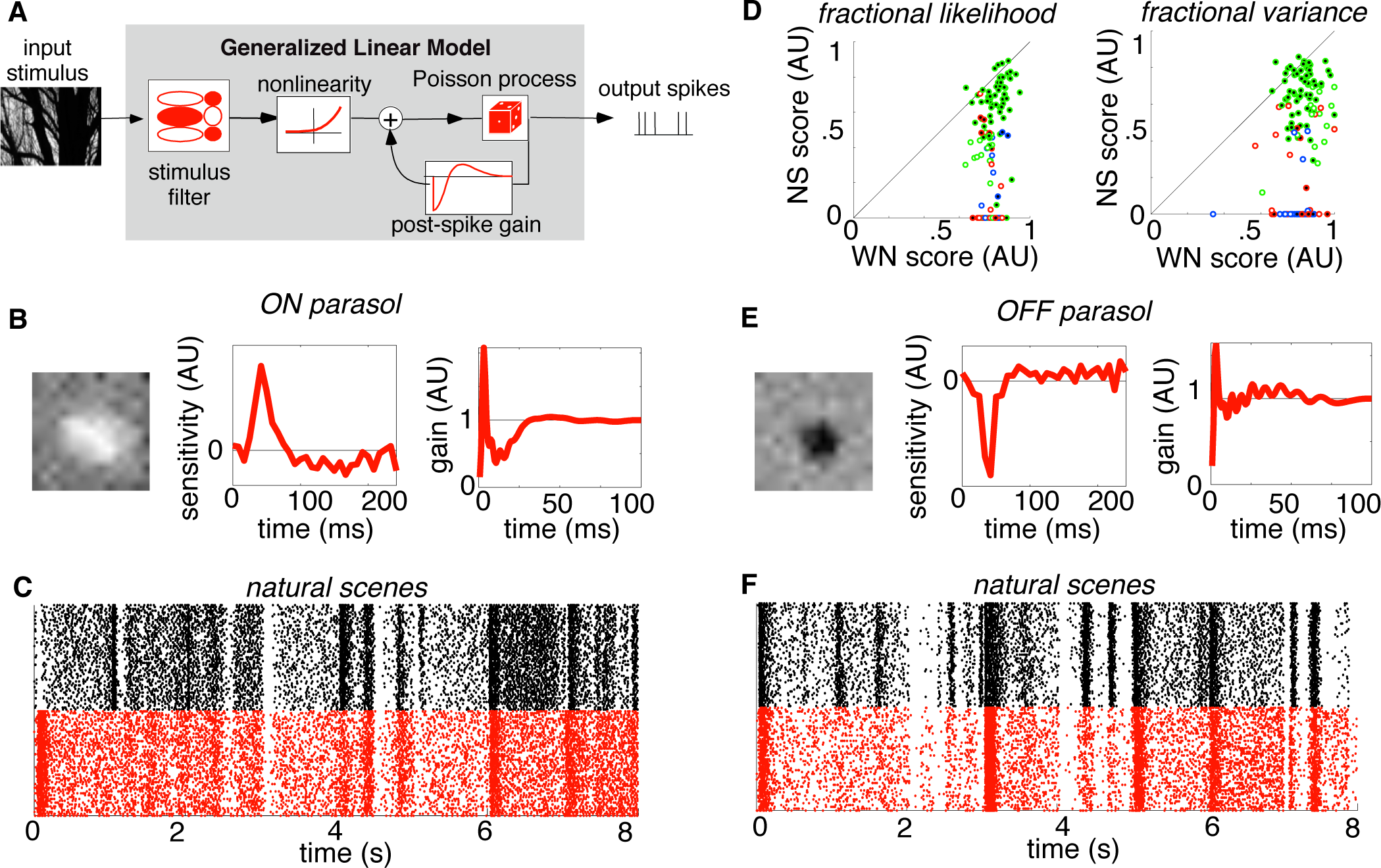
GLM fitted to natural scenes data also fails to predict responses. Details as in Fig. 2. A. Schematic of the GLM. B. Sample GLM components obtained from fits to natural scenes data for an ON parasol cell (same cell as Fig. 2). C. Raster of responses to natural scenes for the same cell (black, 57 repetitions). Response structure is only partially reproduced by the GLM (red). D. Comparison of GLM predictions of natural scenes and white noise responses. In each case the GLM was fitted to the same type of stimulus, cross-validated. E-F. Same as B-C for a sample OFF parasol cell (same cell as Fig. 2).

### Extensions of the model to capture simple nonlinearities

Although the GLM fails to capture responses to natural scenes, it remains possible that slight modifications to the model structure would improve its predictions and thus produce an accurate model, while retaining the technical and conceptual advantages of the GLM, notably the key early linear filtering stage of the model. One modification of this kind is a scalar nonlinearity, that is, a function that transforms a single scalar input to a single scalar output. Such a nonlinearity could capture the function of some nonlinear mechanisms in the retinal circuitry, while only adding a small number of parameters. A nonlinearity of this form can be fitted to data by alternating the relatively simple adjustment of the new parameters with the adjustment of the more numerous GLM parameters that are mathematically guaranteed to converge to a global optimum [24] (see Methods). To test the extensibility of the GLM, the model was augmented to accommodate two pointwise scalar nonlinearities, one before and one after the spatiotemporal linear filter.

First, an instantaneous, scalar nonlinearity was applied to each pixel of the stimulus after temporal integration that would be expected from cone photoreceptors, but prior to the spatiotemporal filter in the GLM (see Methods; Fig 4A). This was performed by convolving the stimulus over time with a realistic photoreceptor impulse response (see Methods) and then fitting the remaining model. This model structure would capture, for example, a nonlinear contrast-response relationship at the cone synapse prior to spatial and temporal processing in the retina. For simplicity, a power-law nonlinear form, f(x) = x^p^, was selected. The parameter governing its shape (p) was iteratively modified, alternating with modifying the parameters of the rest of the model, to maximize likelihood. In three preparations, the form of the fitted nonlinearity was consistent for natural scenes, and consistent for white noise (Fig. 4B, 4C, 4F, 4G), but different for the two types of stimuli. Thus, the early nonlinearity in the model apparently captured modest systematic departures from linearity early in the retinal circuitry, and these departures depend on stimulus statistics. Note that this manipulation would not account for dynamic nonlinearities in photoreceptor signals (e.g. [36]).

**Figure 4.**
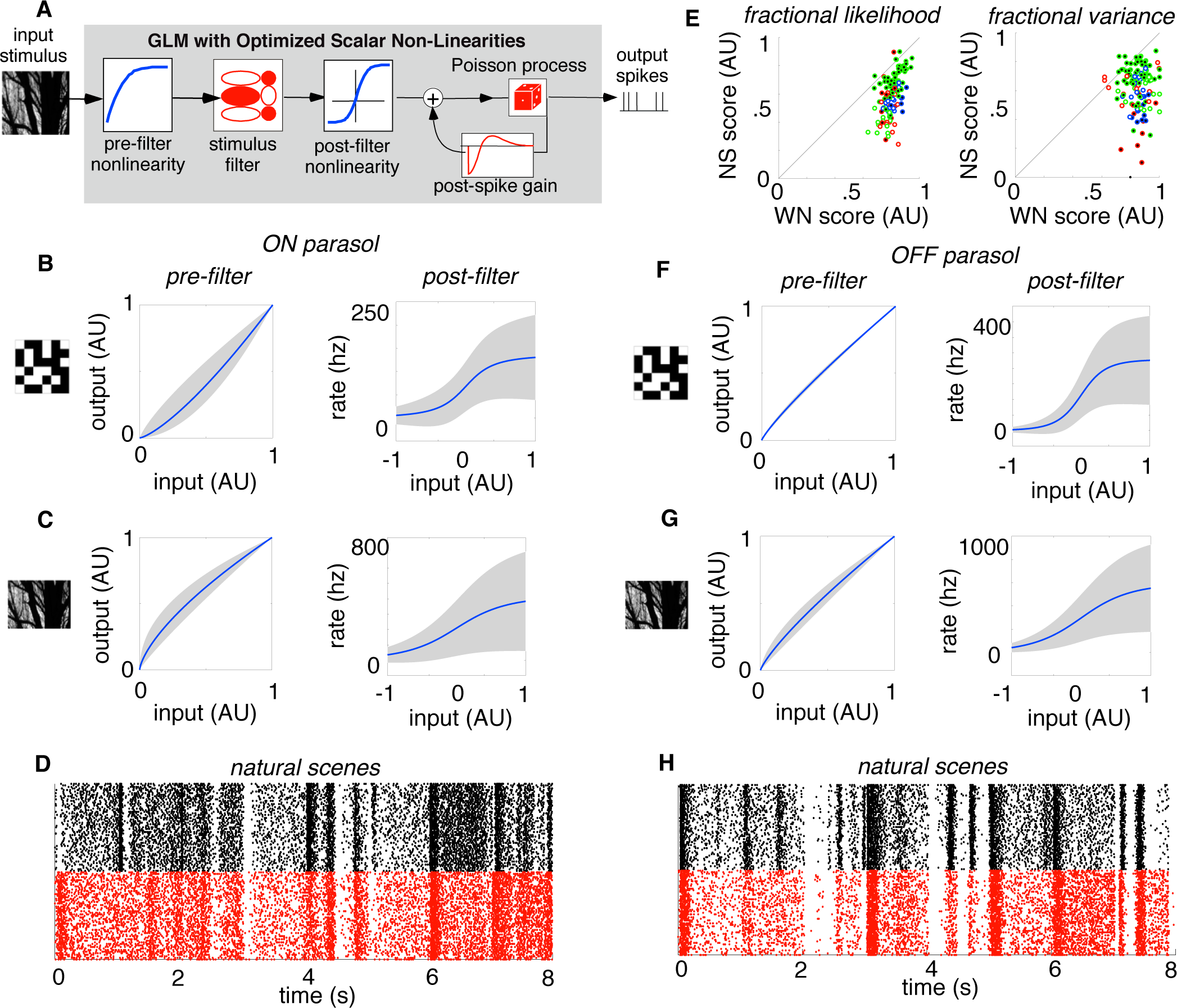
GLM with added scalar nonlinearities performs similarly. Details as in Fig. 3. A. Schematic of the modified GLM, with added nonlinearities highlighted in blue (see text for details). B. Nonlinearities for ON parasol cells fitted with white noise data, with 1 SD variability range across cells given by the gray zone. C. Same as B, except fitted with natural scenes data. D. Raster of responses to natural scenes for an ON parasol cell (same cell as in Fig. 2,3). Response structure is only partially reproduced by the extension of the GLM. E. Comparison of modified GLM predictions of white noise and natural scenes data. F-H. Same as B-D for a sample OFF parasol cell (same cell as in Fig. 2,3).

Second, the exponential nonlinearity in the GLM was modified to allow for different nonlinear transformations of the generator signal controlling firing probability. This model structure would capture, for example, saturation of the RGC firing rate for strong stimuli. A sigmoidal nonlinearity, f(x) = L/(1+exp(-rx)), was fitted by optimizing the parameters governing its shape (L,r), and then refitting the parameters for the feedback filter and the background firing rate, while holding the parameters for the stimulus filter fixed (see Methods). This procedure resulted in reproducible fits with a consistent form across data sets (Fig. 4B,C,F,G).

Fitting both of these simple nonlinearities to the data simultaneously produced some improvement in the natural scenes predictions, as would be expected with a more general model. However, the model continued to produce substantially less accurate predictions with natural scenes than with white noise. Simulated responses reproduced measured responses more accurately than those of the standard GLM, but still featured clearly visible failures (Fig. 4D,H), consistently across cells and data sets (Fig. 4E). In summary, allowing for simple nonlinearities both before and after spatiotemporal linear filtering produced evidence of systematic nonlinear behavior not captured by the original GLM, but did not allow the model to capture responses to natural scenes with substantially higher accuracy. The failure of the modified GLM to capture the response to naturalistic stimuli is thus likely to be attributable to its pseudo-linear structure. Furthermore, the fact that pointwise nonlinearities did not correct the failures implies that more complex nonlinear interactions over space and/or time govern the RGC light response.

## Discussion

The present results show that the generalized linear model (GLM), which successfully captures the responses of populations of primate retinal ganglion cells (RGCs) to white noise stimuli, fails to capture responses as accurately with natural scenes stimuli (Fig. 2). This failure not only reflects the inability of the model parameters to generalize across stimuli with different statistics, but also reflects structural deficits in the model (Fig. 3). Furthermore, simple nonlinearities added to the model to account for known retinal mechanisms had little effect on the conclusions (Fig. 4). The results suggest that models with fundamentally nonlinear structure, potentially including subunits and adaptation, will be necessary to capture how the primate retina represents the visual environment for which it evolved.

### Comparison to expected results from previous models

Since the first experiments demonstrating the center-surround properties of mammalian RGCs [1], many studies have explored the neural computations in the retina which produce patterns of activity transmitted to the brain. Most models of these computations are designed to capture the results of a particular stimulus manipulation (see [17,37]), and provide insight into retinal mechanisms, but are not intended to provide a broad description of the retinal output signal. However, certain models are intended to generalize at least to some degree. These usually have mostly linear properties, so they can be readily interpreted and fitted to diverse experimental data (e.g. [3,4,6,10]). The GLM is such a model: essentially linear, but with simple nonlinear components and conditioning terms added to account for major features of RGC response not captured by simpler models (nonlinear dependence on stimulus strength, non-Poisson time structure such as bursting and refractoriness, and concerted firing across cells). The structure of the GLM allows it to capture a wide variety of response patterns elicited by white noise stimuli, in complete populations of neurons, with striking accuracy [5]. Given its mostly linear structure, however, it is not surprising that substantial failures of the GLM can be observed in some conditions (e.g. Fig. 2D,H) (e.g. [11,13]). It is also not surprising that these failures are larger with natural scenes than with white noise (e.g. Fig. 2C,D,G,H), because white noise only modestly modulates RGC firing and thus may keep responses in a more linear operating range, while natural scenes produce large changes in firing during and between presentation of distinct images.

On the other hand, previous work also suggests that the results presented here could have been quite different. Some studies point to nearly linear signaling in RGCs of some types in some species and conditions (e.g. [10,21,38–41]). Furthermore, nonlinearities previously documented were typically observed with targeted artificial stimuli (e.g. [11,13]), and it is unclear how important these nonlinearities would be for encoding of natural scenes by the retina (but see [42]). Furthermore, early strictly linear models [10] failed to account for important features of light responses that are captured by the GLM, such as accelerating response to increasing stimulus strength, and non-Poisson spike train structure. Therefore, one might have expected the GLM to succeed for a larger range of stimuli, potentially including natural scenes, than the strictly linear models did. Finally, recent work suggested that pseudo-linear models similar to the GLM can capture aspects of RGC responses in the mouse retina to natural scenes [22].

### Simple extensions of the generalized linear model

The utility of models like the GLM is in their capacity to predict the function of the neural circuitry being modeled, without attempting to incorporate all its component parts. In principle, more complex models that capture retinal mechanisms in greater detail would out-perform a simple model like the GLM (e.g. [42]). However, such models typically greatly increase both the difficulty of the fitting process, particularly with complex stimuli such as natural scenes, and the amount of data necessary to constrain the parameters. An alternative approach is to augment the GLM with simple components that can capture a greater range of responses.

The addition of scalar nonlinearities before and after the key spatiotemporal linear step of the GLM (Fig. 4) provided an opportunity to improve its predictive power without substantially increasing the complexity of the model or the fitting procedure, and perhaps to account for significant physiological mechanisms. For example, a scalar nonlinearity could in principle capture nonlinear features of the cone response. In practice, the inferred nonlinearity was modest (Fig. 4B,C,F,G, pre-filter). Also, the exponential nonlinearity in the GLM is highly unrealistic for stimuli that strongly drive firing, while a saturating function is arguably more realistic. Such a function (Fig. 4B,C,F,G, post-filter) produced consistent fits across data sets and cell types, and an improvement in predictions. With these simple nonlinearities, model predictions for natural scenes continued to be substantially inferior to predictions for white noise (Fig. 4), suggesting that the failures of the GLM are more fundamental. However, it is worth noting that different nonlinearities or more flexible fitting procedures could have an impact on the results.

The full GLM has the ability to account for responses of complete populations of cells, including their correlated firing, by the addition of coupling filters [5] that capture interactions between cells. Although the present work was focused on individual cell responses, rather than populations, model fits were also performed with coupling filters to test whether coupling could have an impact on the results. This manipulation had no significant effect (not shown).

### Implications for future work

The present results reveal that neither the GLM nor simple variants of it will capture responses to natural scenes in the primate retina with the fidelity that the GLM exhibited in capturing responses to white noise. This strongly suggests that at least some of the phenomena and mechanisms revealed in many experiments with artificial stimuli (see [17]) will be necessary to explain the output of the retina in natural viewing conditions. Important examples include nonlinear subunits within the receptive field [11], adaptation to stimulus intensity and variance (see [13,14,35,43,44]), and periphery effects [15,45–49]. Future efforts will be needed to determine which aspects of the circuitry in the primate retina are essential for understanding its function in natural conditions.

## Methods

### Recordings

Preparation and recording methods are described elsewhere [29–31]. Briefly, eyes were enucleated from three terminally anesthetized macaque monkeys (Macaca sp.) used by other experimenters in accordance with institutional guidelines for the care and use of animals. Immediately after enucleation, the anterior portion of the eye and the vitreous were removed in room light. Segments of isolated peripheral retina (in Fig. 2-4, red dots 9mm, green dots 7mm, blue dots 11mm temporal equivalent eccentricity [31]) were placed flat, RGC side down, on a planar array of extracellular microelectrodes.

The arrays consisted of 512 electrodes in an isosceles triangular lattice. The arrays had 60 μm inter-electrode spacing and covered a rectangular region 1800 μm by 900 μm. While recording, the retina was perfused with Ames’ solution (34–37°C) bubbled with 95% O_2_ and 5% CO_2_, pH 7.4. Voltage signals on each electrode were bandpass filtered, amplified, and digitized at 20 kHz. A standard spike-sorting algorithm was used to identify spikes from different cells [29] (see below). Only identified cells which exhibited a 1 ms refractory period and plausible electrical image structure were used in further analysis [29].

### Data selection

This study focused on ON parasol and OFF parasol cell types, identified by their characteristic light response properties measured via reverse correlation with white noise stimuli (see [31,32]).

To guarantee that model failures were attributable to the model rather than poor fitting, any cells not meeting a convergence criterion were eliminated from analysis. For each cell, the improvement of the GLM model fit, relative to model of constant firing rate, was calculated with the first half of the fitting data, and then with all of the fitting data. If the improvement over the constant rate model obtained with the first half of the data did not exceed 90% of the improvement obtained with all of the data, then the cell was not analyzed further. This criterion eliminated 221 out of 537 cells.

A second criterion was used to eliminate cells demonstrating unstable light responses. A time-varying rate for each trial of a repeated white noise stimulation was obtained by smoothing the spike train with a Gaussian temporal filter (SD = 10 ms). These data were split into odd and even trials, and the time-varying firing rates in each group of trials were averaged. The odd trial rate was then used as an estimator of the even trial rate, and the fraction of explained variance was computed:

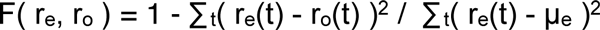

where r_o_(t) is the odd trial rate over time, r_e_(t) is the even trial firing rate over time, and μ_e_ is the average even trial rate. Cells with a value of F below 0.5 were not analyzed further. The same computation was made for the repeated natural scenes presentations, and again cells with a score below 0.5 were eliminated. This criterion eliminated an additional 56 cells.

Finally, all remaining cells were manually screened for spike sorting accuracy. Clusters of recorded spike waveforms in a 3-dimensional space (obtained via principal components analysis) were examined. If the spikes from an identified cell did not form a discrete cluster in both the white noise and natural scenes data, the cell was not analyzed further. This criterion eliminated an additional 127 cells.

### Visual stimulation

Visual stimuli were delivered using the optically reduced image of a CRT monitor refreshing at 120 Hz and focused on the photoreceptor outer segments. The optical path passed through the mostly-transparent electrode array and the retina. The relative emission spectrum of each display primary was measured with a spectroradiometer (PR-701, PhotoResearch) after passing through the optical elements between the display and the retina. The total power of each display primary was measured with a calibrated photodiode (UDT Instruments). The mean photoisomerization rates for the L, M, and S cones were estimated by computing the inner product of the primary power spectra with the spectral sensitivity of each cone type, and multiplying by the effective collecting area of primate cones (0.6 μm^2^ [50,51]). During white noise stimulation, the mean background illumination level resulted in photoisomerization rates of (4100,3900,1600) for the (L,M,S) cones. The naturalistic stimulus resulted in mean photoisomerization rates of (1900,1800,700) for the (L,M,S) cones. Typical visual stimulation lasted roughly a total of 240 min, and initial testing at the same mean light level prior to visual stimulus delivery lasted roughly 30 min. Examination of a fourth data set for which the mean natural scenes intensity was the same as the mean white noise intensity yielded similar results (not shown).

### Generalized linear model (GLM)

The GLM [5] consists of the following stages in sequence: linear spatiotemporal filtering of the visual input, exponential nonlinearity, Poisson spike generation, and a feedback waveform for each spike generated that sums with the post-filter signal (Fig. 2A). All parameters were fitted by maximizing the likelihood of observing the fitting data given the model parameters. The objective function is convex in parameter space; moreover, the resulting gradient and Hessian terms have explicit closed forms [24]. These attributes enable fast and reliable optimization.

The fitted parameters of the GLM consisted of the following: a tonic rate term (μ), a spatiotemporal stimulus filter (k), and a post-spike filter (h). At each time bin of the computed model response, the stimulus filter and post-spike filter contribute to the generator signal by the inner product with the most recent 250 milliseconds of the stimulus (x) and the most recent 100 milliseconds of spiking history (y), respectively. The final instantaneous rate (*λ*) is then determined by exponentiating the sum of the generator signal and the tonic drive term, *λ* = exp(μ + k•x + h•y). Spikes are subsequently generated by a Poisson process with this instantaneous rate.

The stimulus filter maps the high-dimensional spatiotemporal stimulus to a scalar function of time. The filter was constructed by computing the outer product of a spatial filter (169 parameters, one for each location on a 13x13 grid) with a temporal filter (30 parameters, one for each time point over which integration occurs). This produced a spatiotemporal filter that was separable in space and time (rank 1). The spatial filter was centered on the receptive field of the cell estimated from reverse correlation with white noise. Expanding the grid to 15x15 did not improve model performance. The time filter captures the temporal integration of the cell over 250 ms with 8.33 ms samples.

To test the adequacy of the separable filter, for a group of 30 cells an additional pair of space and time filters were fitted, producing a spatiotemporal filter that was the sum of two separable filters (rank 2). This did not improve the natural scenes model performance.

The post-spike filter was constructed from a basis of 20 raised sinusoids [5]. The post-spike filter was further constrained because an unconstrained filter in some cases generated positive feedback and physiologically implausible runaway spiking. To prevent this, the parameter space was restricted so that the time integral of the post-spike filter was no larger than zero.

### Stimulus

The GLM was fitted to both a novel naturalistic stimulus, and to a binary white noise stimulus (96% contrast), both achromatic (L,M and S cones modulated together). The naturalistic stimulus consisted of 900 achromatic images from a standard database [33]. Each image was flipped horizontally and vertically to obtain a total of 3600 unique images. Each image was displayed for one second, and jittered over time according to measurements of fixational eye movements obtained from fixating, awake macaque monkeys (Z.M. Hafed and R.J. Krauzlis, personal communication).

Both types of stimuli were presented in sequences which interleaved the fitting stimulus, consisting of many distinct spatiotemporal sequences, with a test stimulus, consisting of multiple repeats of a shorter spatiotemporal sequence. In two data sets (Fig. 2-4, green dots and blue dots), the naturalistic fitting stimulus was delivered in 60 second segments, with each segment followed by a 30 second presentation of the test stimulus. This sequence of fitting and test stimuli was iterated 59 times. The first two iterations were not used for model fitting and testing to avoid initial transients and adaptation. The white noise fitting stimulus was displayed in 30 second segments, with each such segment followed by a fixed 10 second segment of the test white noise stimulus. This sequence was repeated 60 times. The first three iterations were not used. In the third data set (Fig. 2-4, red dots), the naturalistic fitting stimulus was delivered in 120 second segments with each segment followed by a 60 second presentation of the test stimulus. This sequence was repeated 29 times. The first two iterations were not used. The white noise stimulus for the third dataset (red dots) was identical to the first two datasets (blue and green dots). The overall duration of fitting and testing was approximately the same (~130 minutes) in all three preparations.

### Model evaluation

Two metrics were used in comparing the accuracy of models of light response using white noise and natural scenes stimuli. The first metric, *fractional log-likelihood increment*, is a monotonic transformation of the log-likelihood of the observed data given the model, L(data,model) = log(P(data|model)). A transformation was imposed because likelihood is not a meaningful way to compare responses of a given cell to visual stimuli with different statistics. The focus of this paper is to measure how accurately pseudo-linear models capture fluctuations in light response over time, while accounting for the spiking statistics of RGCs. To achieve this, the fractional log-likelihood increment K was defined as:

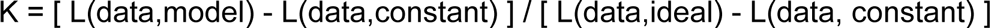

where L(data,constant) is the log-likelihood of the data under a model of constant firing probability over time, and L(data,ideal) is the log-likelihood of the data with an “ideal” model. The ideal model provides an approximate upper bound on the performance of any model which, in the manner of the GLM, predicts smooth response fluctuations over time, and uses spike history feedback to produce realistic spike trains. The stimulus drive r(t) in the ideal model is approximated by computing the empirical time-varying firing rate of the cell in response to repeated presentations of test stimuli, smoothed with a Gaussian temporal filter (SD = 10 ms). Along with this empirical stimulus drive, a post-spike filter and a constant offset term are combined to produce an instantaneous rate, *λ* = exp(μ + c log(r) + h•y), where c is a scalar and h, y, and μ are as given in the GLM above. These parameters of the ideal model are fitted directly to the test data by maximizing likelihood. Note that because both the numerator and denominator of K can be interpreted as bits of information about the stimulus transmitted per spike [52], the ratio can be interpreted as the fraction of the information transmitted by the ideal model. Using this normalization approximately limited the value of K to the range [0,1].

The second metric, *fraction of explainable variance*, compares the recorded and simulated responses to repeated presentations of the test stimulus in a similar manner. For each cell, simulated responses were generated using the model. An average firing rate over time was then calculated across repeats, for both the recorded data and the simulation, and smoothed with a Gaussian temporal filter (SD = 10 ms). Using the simulated rate as a predictor of the recorded rate, the fraction of *explained* variance was computed:

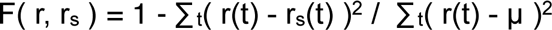

where r(t) is the recorded firing rate over time, r_s_(t) is the simulated firing rate over time, and μ is the average recorded rate. However, this measure does not account for intrinsic variability (noise) in the response of the cell across trials, which cannot be captured by a model that takes as its input the same stimulus on every trial. The measure was therefore normalized by the reproducibility in the recorded repeat responses:

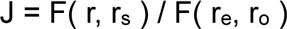

where r_e_ and r_o_ are the even and odd trial firing rates from the data. This results in a metric J indicating the fraction of *explainable* variance, approximately bounded in the range [0,1].

### Input nonlinearity

A possible limitation of the initial linear step in the GLM is that the cone phototransduction process is not fully linear (see [36]). A scalar pointwise nonlinearity was fitted to the data to take this into account. The stimulus sequence was convolved with a temporal kernel which was derived from an estimate of the linear cone impulse response (J.M. Angueyra and F. Rieke, personal communication). Following the temporal kernel, the stimulus intensity values were rescaled to the range [0,1]. A nonlinear transformation was then applied, f(x) = x^p^. The output of this step was used as the input to the GLM. This parameterization permitted smooth exploration of varying degrees of accelerating and compressive nonlinearity. The parameters of the GLM and the nonlinearity were fitted to maximize likelihood in alternating steps. This parameter fitting converged within three iterations.

### Output nonlinearity

A possible limitation of the GLM is the exponential form of the nonlinearity, which is unrealistic for high values of stimulus drive. To deal with this limitation, the exponential was replaced with a sigmoidal function: f(x) = L / (1 + exp(-rx)). The full optimization consisted of four steps. First, the original GLM was fitted. Second, the input to the nonlinearity (x) was defined as the output of the fitted linear filter in the GLM. Third, the two parameters (L,r) of nonlinearity were adjusted to maximize the likelihood of the data, with the other model parameters unchanged. Finally the post-spike filter and tonic drive terms of the GLM (μ,h) were refitted to account for changes induced by the fitted nonlinearity. Following the refitting of μ and h, a second round of fitting L and r produced no increase in likelihood.

## Acknowledgements

This work was supported by National Institutes of Health Grant EY017992 (EJC) and P30 EY019005, and the National Science Foundation Grant 0801700 and DGE-114747 (NB). We thank Clare Hulse, Daniel Ahn, Fumitaka Osakada, and Devon Sandel for technical assistance; Mike Taffe, Tom Albright, and Kristy Berry for access to primate retinas; Matthew Grivich for software development; Steve Barry for machining; Ziad Hafed and Rich Krauzlis for eye movement data; Juan Angueyra and Fred Rieke for cone data. We also thank Fred Rieke, Max Turner, Eero Simoncelli, Jonathan Pillow, and Greg Field for useful conversations and comments on the manuscript.

